# Complexity And Ergodicity In Chaos Game Representation Of Genomic Sequences

**DOI:** 10.1101/2023.12.30.573653

**Authors:** Adel Mehrpooya, Iman Tavassoly

## Abstract

The Chaos Game Representation (CGR) serves as a powerful graphical tool for transforming long one-dimensional sequences, such as genomic sequences, into visually insightful and complex graphical patterns. This algorithmic approach has revealed captivating fractal properties, providing a systems-level perspective on the underlying patterns inherent in specific genomic sequences. Each CGR image generated is distinctive and unique, capturing the individuality of different genomic sequences. We have investigated the nature of the fractal patterns discerned within CGR of genomic sequences, aiming to assess the ergodicity and identify potential ergodic patterns. Our findings present a mathematical proof establishing that CGRs are non-ergodic. This finding suggests that the distribution properties of nucleotide sequences play a pivotal role in shaping the emergent patterns within the CGR images.

## 1. Introduction

Biological systems exhibit a high level of complexity with emergent characteristics evolving over time and space. To comprehend the intricacies encoded within these systems, genomic signal processing emerges as a crucial discipline. It treats genomic sequences as information signals, aiming to find meaningful patterns inherent in the genetic information [1, 2]. In orchestrating cellular processes, especially concerning health and disease, genomic signatures (distinct sets of genes) have been meticulously extracted. These signatures offer a novel lens through which to gain profound insights into the dynamic interplay within cellular networks and responses [3, 4]. Despite genomes serving as the fundamental building blocks of biological data in a dynamic manner, considering them solely as biological sequences can yield valuable insights. This static perspective, when coupled with concepts from information geometry, allows researchers to translate genomic sequences into specific, complex representations, revealing hidden patterns. Information geometry methods, with their emphasis on mathematical structures, provide a unique framework for understanding the relationships and structures within genomic data [2, 5, 6]. The exploration of fractal and self-similar characteristics within genomic sequences further enhances our ability to discern recurring patterns and structures, contributing to a more comprehensive understanding of the underlying biological processes. This holistic and systems-level perspective facilitates the discovery of hidden patterns by exploring the complex layers of information embedded in genomic sequences [7, 8].

Various methodologies have been employed to investigate fractal patterns within genomic sequences. These include random walks, multifractal analysis, Chaos Game Representation (CGR), run analysis, and the assessment of entropy through information theory methods [7–13]. The chaos game is a mathematical technique employed to generate fractal patterns within geometry. The process involves the iterative placement of points within a specified geometric shape, such as a triangle or square, guided by random choices and probabilities. With the systematic addition of points following a predefined set of rules, the result is the formation of self-replicating structures [14, 15]. CGR is an algorithm derived from the chaos game algorithm. It transforms a DNA sequence into a visual representation that exhibits personalized characteristics unique to that specific sequence. CGR has found application in the analysis of protein sequences as well, and various modified versions of the algorithm have been developed for enhanced analytical capabilities. CGRs have been harnessed and integrated into algorithm development, pattern recognition, disease research, phylogenetic studies, as well as functional and comparative genomics [11, 15–21]. CGRs also have inspired the development of machine learning algorithms [22].

Ergodicity refers to the property of a system wherein, over time, the average behavior of the system, as observed through a single trajectory, becomes representative of the behavior averaged over all possible initial conditions [23]. In simpler terms, an ergodic system explores all accessible states and, in the long run, provides a representative sample of its entire state space [24–27]. In the analysis of biological sequences like DNA or protein sequences, ergodic theory facilitates the exploration of the distribution of elements (nucleotides or amino acids) and the identification of patterns that may have functional or structural significance. Given that biological sequences are products of evolutionary processes, ergodic theory aids in modeling the evolutionary dynamics of sequences. This includes understanding how sequences evolve over time, the emergence of specific motifs, and the impact of mutations on sequence patterns [28, 29].

Information theory and CGRs are interconnected in the quest to decode the complex language written in the DNA of living organisms. Genomic sequences, representing the fundamental code of life, embody an enormous amount of information. Information theory provides a conceptual framework to quantify and analyze the information content within these sequences. CGR, on the other hand, detects hidden structures and recurrent motifs within a visual representation of nucleotide sequences. By employing information geometry methods, we can translate genomic sequences into complex landscapes, elucidating the underlying information patterns. This synergy between information theory and CGR offers a unique modality through which we can explore the self-similarities, fractal dimensions, and other emergent features encoded in genomic or any other type of biological sequences [6, 22].

The distribution of nucleotides within a specific genome is not uniform; rather, different genomes exhibit distinct, individualized patterns. In the context of DNA sequences, four nucleotides adenine (A), thymine (T), cytosine (C), and guanine (G)are represented by single letters. The ratio of adenine to thymine (A*/*T) is commonly referred to as the AT content, while the ratio of cytosine to guanine (C*/*G) represents the GC content. These ratios vary across genomes, carrying implications for diverse biological processes and functions. GC skew, measuring the relative abundance of guanine over cytosine (G − C) or vice versa, is indicated by positive values for guanine excess and negative values for cytosine excess. Similarly, AT skew assesses the relative abundance of adenine over thymine (A − T) or vice versa, with positive values denoting adenine excess and negative values indicating thymine excess. Notably, nucleotide distribution is not uniform throughout the entire genome; local variations, such as CpG islands (regions rich in CG dinucleotides), play crucial roles in regulatory processes. In protein-coding regions, the phenomenon of codon usage bias, where synonymous codons (encoding the same amino acid) are employed with varying frequencies, contributes to nucleotide distribution variations. Some genomes feature repetitive elements, including short tandem repeats (microsatellites) and long interspersed nuclear elements (LINEs), which disrupt uniform nucleotide distribution. This non-uniformity extends to different genomic regions, including coding regions (exons), non-coding regions (introns), and intergenic regions. The overall composition of a genome, quantified in terms of the percentages of A, T, C, and G, exhibits wide variation among different organisms, commonly known as genomic base composition. The study of nucleotide distribution patterns is vital for comprehending the functional and structural characteristics of genomes. Techniques such as CGR and various bioinformatics tools serve as valuable aids in visualizing and analyzing these patterns, providing insights into genome organization and evolution [30–32].

## 2. CGR Algorithm

CGR is a method to visually represent DNA sequences in a fractal-like pattern. In the chaos game representation of genomic sequences, each nucleotide (A, C, G, T) is associated with a specific position in a coordinate system [10, 33]. The algorithm proceeds by iteratively plotting points based on the sequence. Here’s a step-by-step algorithm assuming the coordinate system you provided:

1. **Initialize Coordinates:** Set up a coordinate system with four points corresponding to the nucleotides:
  - C (Cytosine) at the left upper corner.
  - G (Guanine) at the right upper corner.
  - A (Adenine) at the left lower corner.
  - T (Thymine) at the right lower corner.
2. **Choose a Starting Point:** Select a starting point (can be arbitrary) within the coordinate system.
3. **Iterate through the Genomic Sequence:** For each nucleotide in the genomic sequence:
  a. Calculate the midpoint between the current point and the point associated with the nucleotide.
  b. Move to the calculated midpoint.
  c. Plot a point at the new position.
4. **Repeat:** Repeat the process for each nucleotide in the sequence.
5. **Visualization:** After iterating through the entire genomic sequence, you should have a set of plotted points that form a pattern within the coordinate system.

Figure 1 illustrates the fundamental steps involved in generating CGR for any arbitrary DNA sequence.

**Figure 1.**
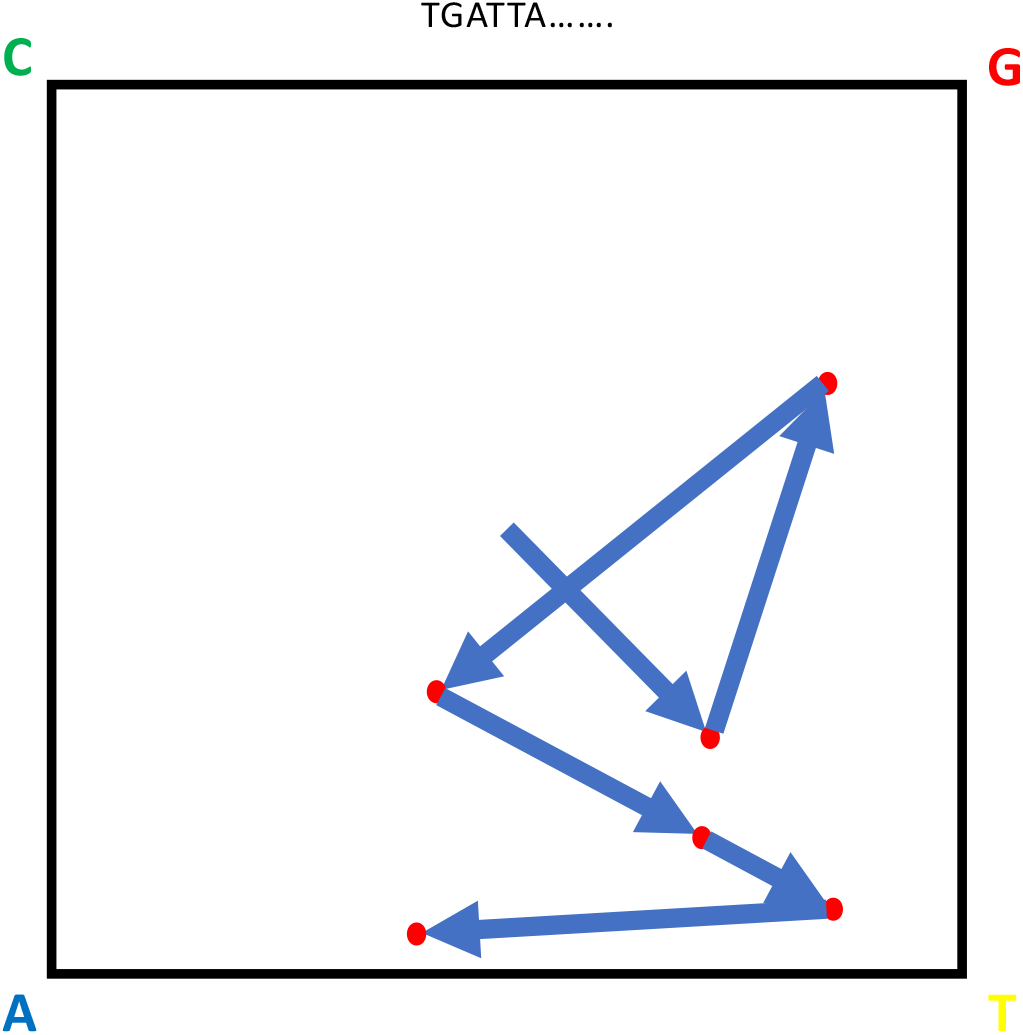
The process of generating a CGR using a genomic sequence.

In Figure 2, CGRs of two genomic sequences are depicted: the human mitochondrial DNA (mtDNA) (NCBI Reference Sequence: NC012920.1) and the complete genome of Human Immunodeficiency Virus type 1 (HIV-1) (NCBI Reference Sequence: AF033819.3).

**Figure 2.**
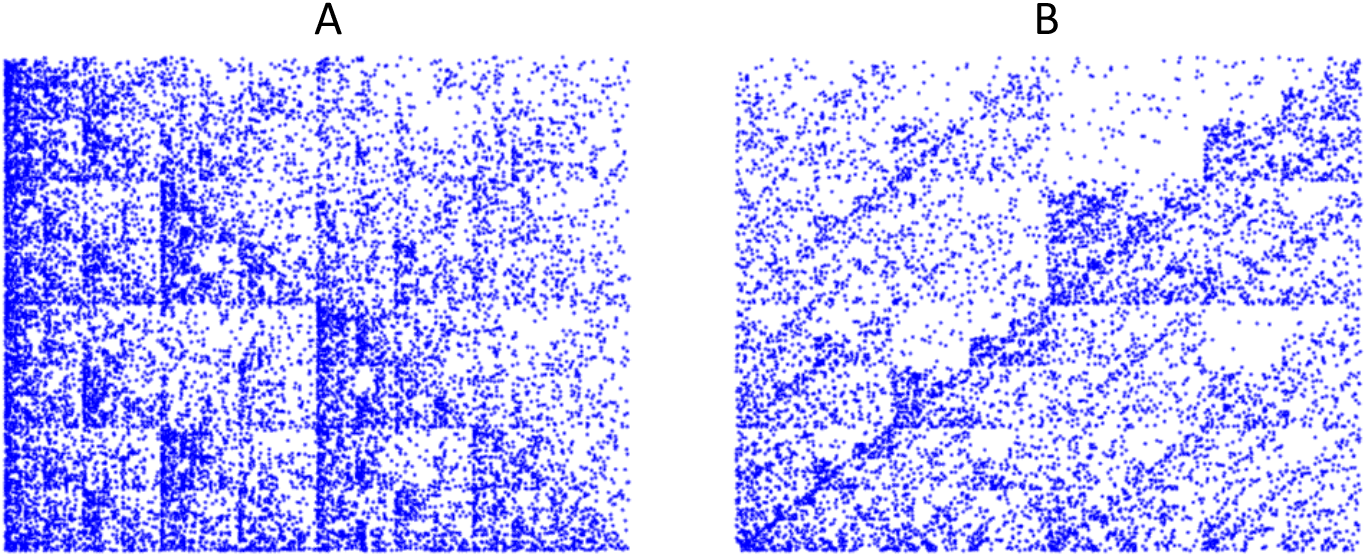
A: CGR of human mtDNA genome. B: CGR of HIV-1 genome.

If we assign four distinct colors to each dot representing the four nucleotides, the CGR will exhibit a division into four distinct regions, each distinguished by its specific color. Figure 3 illustrates this modification in the CGR algorithm applied to the entire genomes of both human mtDNA and HIV-1. Building on our earlier discussion, the patterns that emerge from CGRs are influenced by the content and distribution of nucleotide sequences. Figure 4 showcases both the conventional CGR and a modified version (employing distinct colors for each nucleotide) for a synthetic DNA sequence generated *in silico* through the creation of random sequences. In these CGRs, no discernible pattern emerges; rather, an even distribution of dots is observed across the squares.

**Figure 3.**
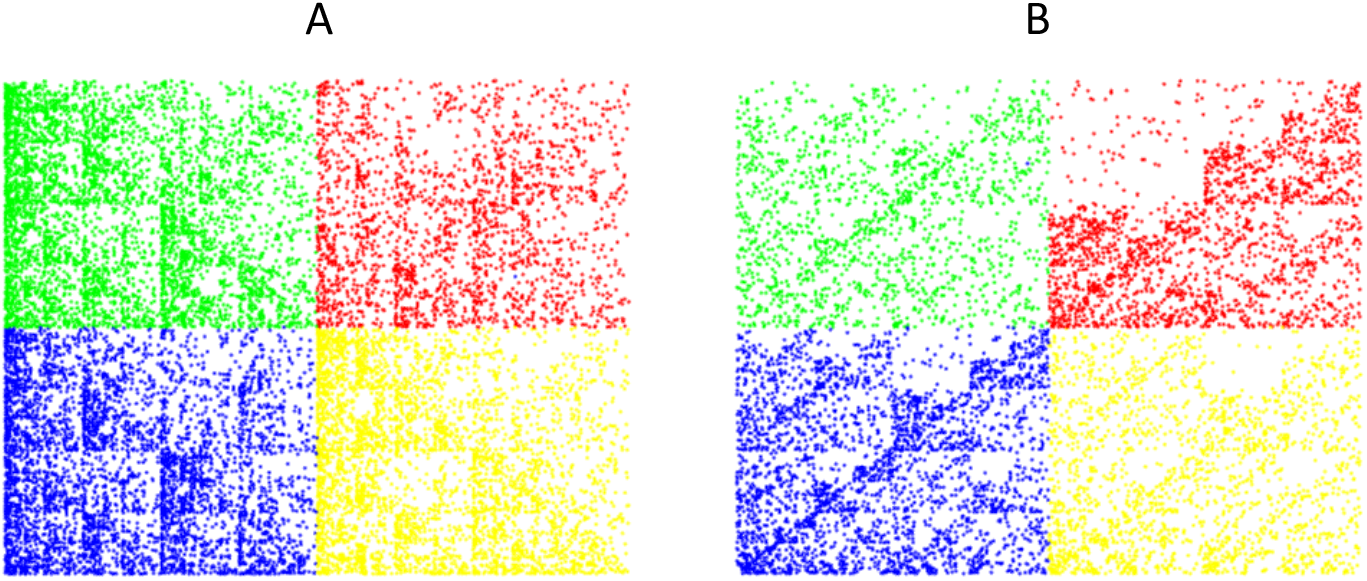
Colorful version of CGR using different colors for each nucleotide: A: Human mtDNA genome. B: HIV-1 genome.

**Figure 4.**
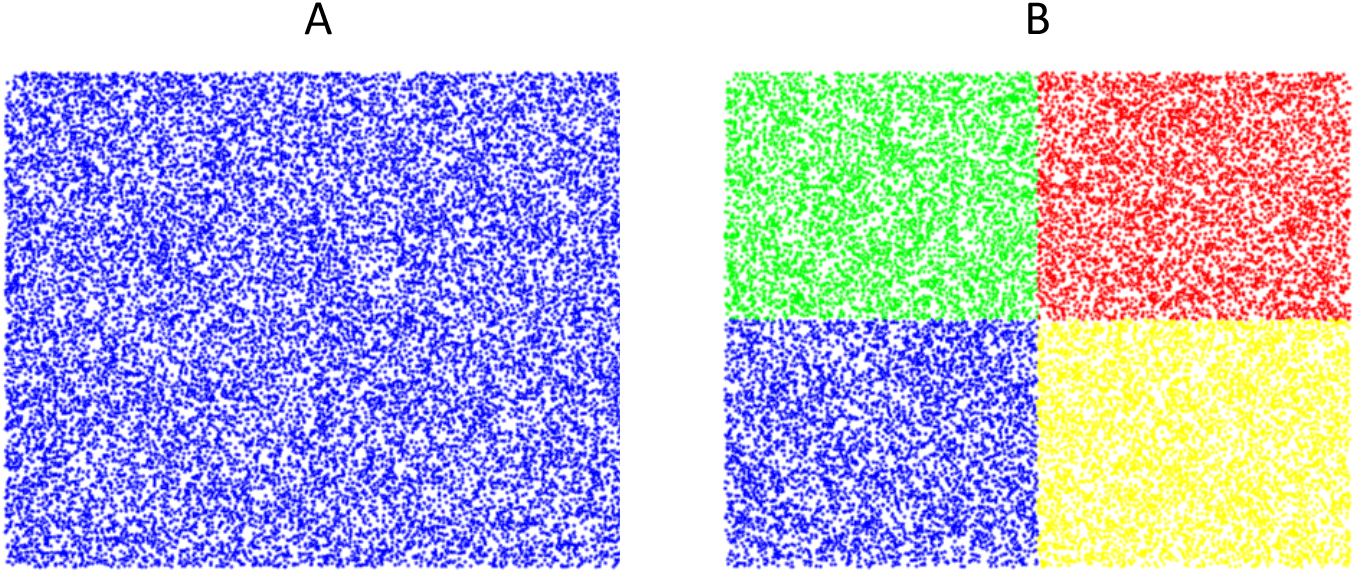
CGR for a randomly generated DNA sequence: A: Regular CGR. B: Colorful version of CGR using different colors for each nucleotide.

Moreover, it is essential to highlight the distinction between the distribution patterns observed in CGRs for real DNA sequences and those generated synthetically through random sequences. Real DNA sequences, being products of biological processes, exhibit inherent patterns and structures reflective of their biological functions. In contrast, the synthetic DNA sequences generated randomly lack the intricate organization inherent in natural DNA. The even distribution of dots in the CGRs of the synthetic sequences contrasts sharply with the characteristic patterns often discerned in CGRs derived from genuine genomic data. This divergence underscores the importance of considering the biological context and underlying structure when interpreting CGRs of DNA sequences. Supplemental Video 1 showcases the production process of Human mtDNA CGR.

## 3. CGR, Ergodicity of genomic Sequences and Distribution of Runs

Here, we present a comprehensive overview of invariant measures for iterated function systems (IFSs), explore the ergodicity of IFSs, and delve into the connections between CGR and IFS ergodicity. Additionally, we consider a statistical approach to analyze genomic sequences, potentially influencing the analysis of CGR images. This method examines the relationship between genomic sequences and the statistical distribution of runs within those sequences.

In 1970, Brunel demonstrated a necessary and sufficient condition for a Markovian operator *P* to possess an invariant measure, stating that each convex combination of iterations 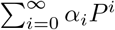 must be conservative [34]. Building upon Brunel’s work, Horowitz in 1972 investigated finite invariant measures for sets of Markov operators and extended Brunel’s result to commutative semigroups of Markovian operators [35]. Further exploration focused on Iterated Function Systems (IFSs) employing regenerative sequences for iterations, establishing ergodic theorems under log-average contractivity conditions [36].

In 2000, a novel criterion was introduced to ascertain the existence of invariant distributions for Markov operators [37]. Subsequently, a survey highlighted crucial results concerning the uniqueness of invariant probability measures for place-dependent random IFSs [38]. Investigating Markov chains generated by IFSs, the study addressed the aperiodic strong ergodicity concerning time series [39]. In 2008, a topological perspective explored invariant sets of IFSs, revealing that the family of all invariant sets is a nowhere dense *F*_*σ*_ subset within a metric space composed of nonempty compact subsets of ℝ^N^ endowed with the Hausdorff metric [40].

Significantly, the projection entropy *h*_*π*_, defined on the code space of an IFS, was established, leading to the study of essential ergodic characteristics related to it. This concept proved instrumental in examining the dimensional characteristics of invariant measures on the IFS attractor and establishing a variational principle between ergodic measures’ projections and the Hausdorff dimension of the attractors [41].

Notably, Oliver et al. (1993) applied entropic profiles in DNA sequences based on CGR. This approach effectively discerns random and natural DNA sequences, highlighting distinct variability levels within and between genomes [42]. Further advancements include conditions discovered in 2010, determining when the chaos game algorithm almost surely yields the IFS attractor [43]. In 2015, the ergodicity of Lebesgue measures for IFSs of diffeomorphisms was explored, providing sufficient conditions for the minimality of IFSs on continuum spaces to ensure Lebesgue measure ergodicity [44].

Targeting specific weakly hyperbolic sequences on IFS code spaces, recent research delved into strict attractor properties and conditions where the chaos game produces target sets, establishing asymptotic stability for IFSs on [0, 1] [45]. Additionally, investigations in 2017 focused on the existence of attractors and the ergodicity of invariant measures for weakly hyperbolic IFSs on compact metric spaces [46].

We commence with a comprehensive review of the ergodicity of IFSs and explore related results. Subsequently, we engage in a concise discussion regarding potential connections between CGR and ergodic theory. Finally, we delve into the investigation of the run count approach.

Let (*X*_1_, *B*_1_, *μ*_1_) and (*X*_2_, *B*_2_, *μ*_2_) represent two probability spaces. A transformation *T* : *X*_1_ ⟶ *X*_2_ is termed measure-preserving if it satisfies the following conditions:

1. *T* is measurable; that is for every *β*^*′*^ ∈ *B*_2_, we have *T* ^*−*1^(*β*^*′*^) ∈ *B*_1_;
2. *T* preserves *μ*_1_ and *μ*_2_; that is for every *β*^*′*^ ∈ *B*_2_, we have *μ*_1_(*T* ^*−*1^(*β*^*′*^)) = *μ*_2_(*β*^*′*^). Additionally, an invertible measure-preserving transformation, *T*, is bijective, and *T* ^*−*1^ is also measure-preserving.

### Proposition. 3.1

The following statements are held.

1. For two measure-preserving transformations *T*_1_ : *X*_1_ ⟶ *X*_2_ and *T*_2_ : *X*_2_ ⟶ *X*_3_, *T*_2_ ? *T*_1_ : *X*_1_ ⟶ *X*_3_ is also measure-preserving.
2. If *T*_*i*_ : (*X*_*i*_, *B*_*i*_, *μ*_*i*_) ⟶ (*X*_*i*_, *B*_*i*_, *μ*_*i*_) are measure-preserving, where *i* = 1, 2, then *T*_1_ × *T*_2_ : (*X*_1_ × *X*_2_, *B*_1_ × *B*_2_, *μ*_1_ × *μ*_2_) ⟶ (*X*_1_ × *X*_2_, *B*_1_ × *B*_2_, *μ*_1_ × *μ*_2_) is measure-preserving.

Since the iterates of *T*, that is *T*^*n*^ for large *n*, is of primary concern in our investigation, in the rest of this section, it is supposed that (*X*_1_, *B*_1_, *μ*_1_) = (*X*_2_, *B*_2_, *μ*_2_).

### Definition. 3.2.

Let *T* be a measure-preserving transformation of the probability space (*X, B, μ*). *T* is said to be ergodic if *T* ^*−*1^(*β*) = *β* implies that *μ*(*β*) = 0 or *μ*(*β*) = 1, where *β* ∈ *B*.

### Theorem. 3.3.

Let *T* be a measure preserving transformation of the probability space (*X, B, μ*). Then statements (1) and (2) are equivalent.

1. *T* is ergodic;
2. if *f* : (*X, B, μ*) ⟶ (*X, B, μ*) is measurable and for every *x* ∈ *X*, we have (*foT* )(*x*) = *f* (*x*), then *f* is constant almost everywhere.

Let *X* be a compact metrizable space. Assume that *d* is a metric on *X* and *B*(*X*) is the *σ*– algebra of Borel subsets of *X*. This means that *B*(*X*) is the smallest *σ*–algebra of *X* containing all open subsets of *X*. Let *P* (*X*) be the collection of all Borel probability measures on (*X, B*(*X*)). For a continuous transformation *T* : *X* ⟶ *X*, a mapping 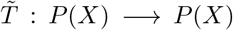 is defined by 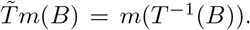. The set 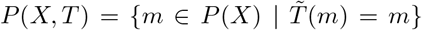 is composed of all Borel measures for which *T* is measure-preserving.

### Theorem. 3.4.

Let *m* ∈ *P* (*X*) for which *m*(*U* ) *>* 0 for every non-empty open set *U* of *X*, and suppose *T* : (*X, B*(*X*), *m*) ⟶ (*X, B*(*X*), *m*) is a continuous and ergodic transformation that preserves *m*. Then almost every orbit of *X* is dense.

### Theorem. 3.5.

Let *T* : *X* ⟶ *X* be a continuous mapping and *m* be a probability measure on the Borel probability measurable space (*X, B*(*X*)). Then *m* ∈ *P* (*X, T* ) if and only if ∫ *foTdm* = ∫ *fdm* for every continuous mapping *f* : *X* ⟶ *X*.

The IFSs can be studied through a similar approach.

### Definition. 3.6.

Let *f* : (*X, d*) ⟶ (*X, d*) be a function on a complete metric space *X*. The mapping *f* is said to be a contraction with the contraction factor *ξ* if for some *c* ∈ [0, 1), we have:

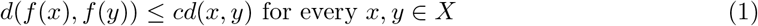

and *ξ* = min{*c* ∈ [0, 1) | *c* stisfies Eq. (1)}. The collection of all contractions on *X* is denoted by Lip_1_(*X*).

Let chaos game be played for four points *A* = (0, 0), *C* = (0, 1), *G* = (1, 1) and *T* = (1, 0). The area *X* that is surrounded by the square whose vertices are the points *A, B, C* and *D*, endowed with Euclidean metric is a compact metric space. Suppose that *p* ∈ *X* is arbitrary. The subspace *P*_1_(*X*) of *P* (*X*) is defined as:

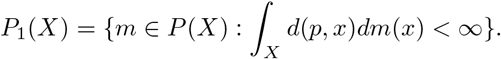

*P*_1_(*X*) is a metric space endowed with the *MK* metric defined by:

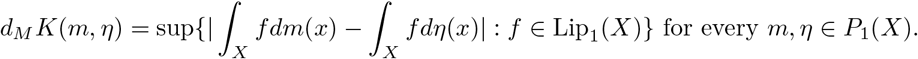

In case the chaos game is played based on iterated function systems with probability, IFSP, the Markov operator can be defined.

### Definition. 3.7.

The Markov operator *M* : *P* (*X*) ⟶ *P* (*X*) associated with an IFSP is given by:

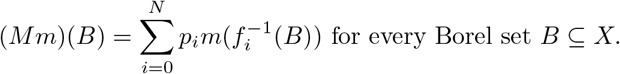

### Definition. 3.8.

Let *X* be a complete and separable metric space. Then (*P*_1_(*X*), *d*_*MK*_) is complete. In case *X* is compact, then (*P* (*X*), *d*_*MK*_) = (*P*_1_(*X*), *d*_*MK*_) and these spaces are also compact.

### Theorem. 3.9.

Let {*f*_*i*_, *p*_*i*_} be an IFSP and suppose that *c*_*i*_ is the contraction factor of *f*_*i*_. Then

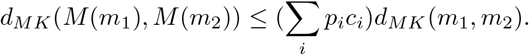

### Corollary. 3.10.

If ∑_*i*_ *p*_*i*_*c*_*i*_ *<* 1, then there exists a unique Borel probability measure *m* ∈ *P*_1_(*X*) such that:

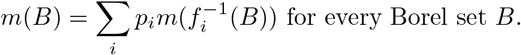

To understand the relationship between the attractor of the IFS *f*_*i*_ and the invariant measure *m* established based on *f*_*i*_ in an Iterated Function System with Probability (IFSP), it is crucial to consider the concept of the support of a measure. The support of a measure *m* ∈ *M* (*X*) is defined as:

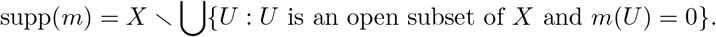

### Theorem. 3.11.

Let {*f*_*i*_, *p*_*i*_} be an IFSP and suppose that *c*_*i*_ is the contraction factor of *f*_*i*_ so that for every *i, p*_*i*_ *>* 0 and *c*_*i*_ *<* 1. Then supp(*m*) = *A*, where *m* is the invariant measure and *A* is the attractor of the IFS {*f*_*i*_}.

### Definition. 3.12.

In the context of an Iterated Function System *f*_*i*_, an invariant set *A* is defined as 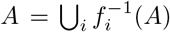. For a probability measure *m* ∈ *P* (*X*) with respect to which *f*_*i*_ is measure-preserving, the IFS *f*_*i*_ is considered ergodic if, for every invariant set *A* within *f*_*i*_, the measure *m*(*A*) satisfies *m*(*A*) ∈ {0, 1}.

It is evident that an IFS, whose maps are derived from a nucleotide sequence, can be transformed into an IFSP. This transformation involves assigning a probability to each map corresponding to the points A, C, G, or T. The probability is defined as the quotient of the frequency of the corresponding letter in the nucleotide sequence and the total sequence length. Hence, every IFS associated with a nucleotide sequence can be seen as an IFSP.

By selecting the IFSP maps for the CGR so that their contraction factors, *c*_*i*_, satisfy Eq. (2),

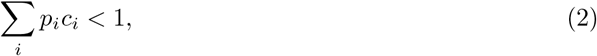

it is possible, as per Corollary 3.10, to determine a measure *m*. This measure is such that the IFSP becomes measure-preserving with respect to it. However, if *A* ∈ *B*(*X*) is an invariant set for the given IFSP, it is important to note that *X* ∖ *A* is not necessarily invariant. This means that decomposing the IFSP mappings into two smaller domains, *A* and *X* ∖ *A*, when the IFSP is not ergodic, is not guaranteed. Consequently, the decomposition of the corresponding CGR image into two disjoint parts, which could simplify the analysis, might not be feasible.

Other problems related to ergodic theory and CGR include the study of entropy in CGR images [41, 42] and invariant measures for Markov chains modeling IFSPs corresponding to DNA sequences [39, 40, 43–46].

Another mathematical approach to genomic sequences involves run distribution analysis in DNA sequences, offering a statistical perspective. This theoretical framework has found success across diverse applications, including nonparametric hypothesis testing, reliability theory, quality control, general applied probability, computer science, and genome analysis. The versatility of run distribution analysis underscores its effectiveness in extracting valuable insights from genomic data, contributing to advancements in various fields [47].

Sprizhitsky et al. conducted a statistical analysis of nucleotide runs in DNA sequences from various species, including vertebrates, prokaryotes, and invertebrates, revealing a considerable variety of run distributions among those species [48]. Louchard employed a Markov chain approach to investigate asymptotic properties of monotone-increasing runs of uniformly distributed integer random variables, applying the findings to DNA analysis [49]. Ponty et al. introduced a methodology and software tool for randomly generating genomic sequences and structures. Their methodology, integrating structural parameters, enhanced classical parametric Markovian models used as frameworks for analyzing occurrences of oligomers in biological sequences. Moreover, their approach proved practical in addressing various classes of models for sequence analysis, including Markov chains, hidden Markov models, weighted context-free grammars, regular expressions, and PROSITE expressions [50].

Nuel et al. tackled the problem of counting occurrences in a set of independent sequences generated by a Markov source. They developed efficient methods and algorithms for such computations in the context of homogeneous or heterogeneous Markov models. These established algorithms were then tested on three biological datasets [51]. Johnson et al., in [47], considered the asymptotic relative error of normal, Poisson, compound Poisson, and finite Markov chain embedding, as well as large deviation approximations regarding approximating the distributions of the number of runs and patterns. Their numerical comparisons helped determine the methods with the best performance for approximating run distributions.

For a given sequence composed of elements from a finite set *S* = {*s*_1_, …, *s*_*k*_}, a run is defined as a sequence of one or more *s*_*i*_, where *i* ∈ {1, …, *k*}. In the context of *N* ordered observations, where *n*_1_ are of one kind and *n*_2_ = *N* − *n*_1_ are of another kind, a test can be established based on the number of runs in a sequence of *N* ordered observations. The hypothesis that the terms of a given sequence are random is rejected if too few or too many runs exist within the sequence [52]. A DNA sequence can be transformed into a sequence of Bernoulli trials, specifically, a sequence composed of two symbols *X*_1_ and *X*_2_, where each nucleotide corresponds to an element of the transformed sequence. For instance, in the analysis of Purine, *X*_1_ represents *A* and *G*, and *X*_2_ represents *T* and *C*.

The investigation focuses on determining the number of runs in this Bernoulli sequence generated through a specific method. Identifying a run of length *n* is seen as a successful outcome within the sequence of Bernoulli trials. Statistical distributions of runs in various DNA sequences can be established, enabling mathematical comparisons of their structural characteristics.

Analyzing the statistical distribution of runs in a DNA sequence involves defining random variables *R*_1_ and *R*_2_. These variables represent the counts of runs, denoted by *X*_1_ and *X*_2_ respectively, in the Bernoulli sequence obtained by transforming the provided DNA sequence. Examining the joint probability distribution function of *R*_1_ and *R*_2_ unveils the interrelation and probability characteristics of runs in the transformed Bernoulli sequence. The joint probability distribution function of *R*_1_ and *R*_2_ is given by:

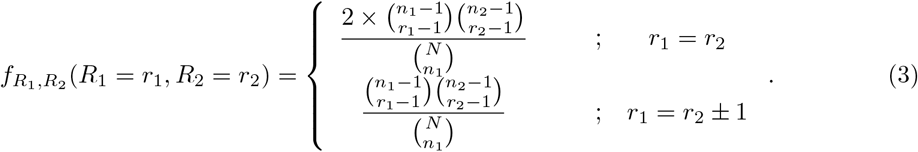

Now, let *R* denote the random variable that counts the number of runs for a given DNA sequence. Considering Eq. (3), the probability distribution function of *R* is given by:

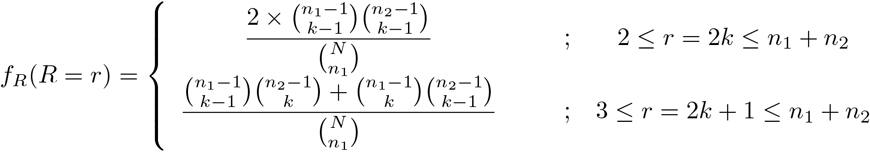

Moreover, the mean and variance of *R* can be calculated by the formulas (4) [52]:

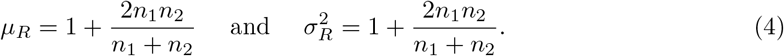

For a notable example of the application of the run count approach, refer to [48], where this method was applied to DNA sequences by comparing the number of observed runs for a DNA sequence with expected values for a random sequence.

## 4. Conclusions

In conclusion, our rigorous mathematical analysis has conclusively demonstrated that CGRs lack ergodic properties. In particular, when the IFSP lacks ergodicity, there is no assurance of successfully decomposing the IFSP mappings into two smaller domains, *A* and *X* ∖ *A*. As a result, the decomposition of the corresponding CGR image into two separate parts, aimed at simplifying the analysis, may not be achievable. This underscores the profound impact of nucleotide sequence distribution in shaping the distinct structures inherent in CGRs. The intricate interplay between the distribution patterns and the resulting CGR structures opens a compelling avenue for further research.

Moving forward, there is an urgent imperative for comprehensive investigations aimed at precisely defining and detecting these distribution models. This is particularly crucial given their potential implications for genome evolution and their relevance to various diseases and health conditions. The sensitivity of CGR structures to genomic aberrations emerges as a pivotal area for exploration, demanding meticulous scrutiny to unveil the underlying mechanisms.

Notably, the non-random nature of genomic sequences prompts a deeper exploration of the complex distributions across different species, health status, and disease conditions. Identifying the influencing parameters in this context holds immense potential, offering a nuanced understanding of the implications and applications of CGRs in the field of systems biomedicine.

This pursuit promises to yield valuable insights that can significantly contribute to advancements in biomedical research and applications. By unraveling the complexities of CGR structures and their relationship to genomic information, we are poised to enhance our understanding of biological systems and pave the way for innovative approaches to diagnosis, treatment, and disease prevention.

## Algorithms and codes

The codes for generating CGR images from specific sequences are conveniently provided as supplemental files, offering versatility across different programming languages. These supplementary materials encompass code implementations in Matlab, Python, and R.

## Supporting information

Supplemental Movie 1-Human mtDNA-CGR Generation Movie

CGR Generation-Matlab Code

CGR Generation-Python Code

CGR Generation-R Code

