## Supplementary material for "Complexity And Ergodicity In Chaos Game Representation Of Genomic Sequences": CGR Generation-Python Code

#This code converts a genomic sequence in FASTA format to a CGR image. 
#Reference: MEHRPOOYA A,  Tavassoly I, 2023.


# Read the genomic sequence in FASTA format
record = next(SeqIO.parse("sequence.fasta", "fasta"))
sequence = str(record.seq)

# Initialize variables
x, y = 50, 50
n = len(sequence)

# Lists to store coordinates
x_values, y_values = [], []

# Iterate through the sequence
for i in range(n):
    if sequence[i] == "C":
        x *= 0.5
        y = (y + 100) * 0.5
    elif sequence[i] == "G":
        x = (x + 100) * 0.5
        y = (y + 100) * 0.5
    elif sequence[i] == "A":
        x *= 0.5
        y *= 0.5
    elif sequence[i] == "T":
        x = (x + 100) * 0.5
        y *= 0.5
    x_values.append(x)
    y_values.append(y)

# Plot the CGR
plt.scatter(x_values, y_values, color='blue')
plt.axis('off')
plt.show()
