## Supplementary material for "Complexity And Ergodicity In Chaos Game Representation Of Genomic Sequences": CGR Generation-R Code

#This code converts a genomic sequence in FASTA format to a CGR image. 
#Reference: MEHRPOOYA A,  Tavassoly I, 2023.

library(seqinr)
library(ggplot2)

# Read the genomic sequence in FASTA format
sequence <- read.fasta(file = "sequence.fasta")

# Initialize variables
x <- 50
y <- 50
n <- nchar(sequence$seq)

# Create a data frame to store coordinates
data <- data.frame(x = numeric(n), y = numeric(n))

# Iterate through the sequence
for (i in 1:n) {
  if (substring(sequence$seq, i, i) == "C") {
    x <- x * 0.5
    y <- (y + 100) * 0.5
  } else if (substring(sequence$seq, i, i) == "G") {
    x <- (x + 100) * 0.5
    y <- (y + 100) * 0.5
  } else if (substring(sequence$seq, i, i) == "A") {
    x <- x * 0.5
    y <- y * 0.5
  } else if (substring(sequence$seq, i, i) == "T") {
    x <- (x + 100) * 0.5
    y <- y * 0.5
  }
  data[i,] <- c(x, y)
}

# Plot the CGR
ggplot(data, aes(x, y)) +
  geom_point(color = "blue") +
  theme_void()
